# Integration of protein and coding sequences enables mutual augmentation of the language model

**DOI:** 10.1101/2024.10.24.620004

**Authors:** Heng-Rui Zhao, Meng-Ting Cheng, Jinhua Zhu, Hao Wang, Xiang-Rui Yang, Bo Wang, Yuan-Xin Sun, Ming-Hao Fang, Enhong Chen, Houqiang Li, Shu-Jing Han, Yuxing Chen, Cong-Zhao Zhou

## Abstract

Recent language models have significantly accelerated our understanding on the massive biological data, using protein or DNA/RNA sequences as a single-language modality. Here we present a dual-language foundation model, which integrates both protein and coding sequences (CDS) for pre-training. Compared to the benchmark models, it shows a superior performance up to ∼20% on both protein and mRNA-related discriminative tasks, and gains the capacity to de novo generate coding sequences of ∼50% increased protein yield. Moreover, the model also possesses the knowledge transferability from the pre-training data to the upstream 5’ untranslated regions. These findings indicate the intrinsic correlations between protein and its CDS, as well as the coding region and beyond. It provides a new paradigm that leverages the multiple-language foundation model to interpret the hidden context of distinct corpora/biological languages, which could be further applied to mine the yet-unknown biological information/correlation beyond the Central Dogma.

## Introduction

The Central Dogma, one of the most fundamental concepts in biology, describes the flow of genetic information within all living organisms, strongly indicating the intrinsic correlation between DNA, mRNA and protein sequences. It is generally accepted that mRNA plays a crucial role in transferring the genetic information from DNA to proteins, driving the precise execution of diverse biological functions. The coding sequences (CDS) of mRNA operates 64 nucleotide triplets, known as codons, each of which encodes a specific amino acid residue, except for the stop codons. Beyond the CDS, the upstream 5′ untranslated region (5′ UTR) of mRNA also modulates the translation initiation efficiency and determines the spatially and temporally specific synthesis of protein via adopting variable configurations^1–3^. The molecular biological investigations in past half a century generally focus on the linear correlation and synteny transfer of information from DNA/RNA sequences and encoded protein sequences in a one-dimensional manner. It results in the loss of vast intrinsic disciplines in the context, such as the correlation between a given nucleotide/codon and another distal one, even the crosstalk between a segment of CDS and an independent protein fragment in the same pair of CDS/protein.

The emerged technologies of natural language processing (NLP) might provide a new paradigm of the biological research to solve these puzzles, given the similarity of DNA/RNA/proteins sequences to the natural languages. In fact, transformer-based models, such as BERT and GPT^4–6^, have achieved remarkable successes in the natural language understanding (NLU) and generation (NLG). Recently, transformer-derived techniques have also been utilized for protein language models (PLM) such as ESM^7^, ProtT5^8^, and Ankh^9^, demonstrating the high effectiveness and adaptability in better understanding of the protein structure and function. For instance, ESM2 leverage its power in semantic and situational compression of protein sequences, enabling the interpretation of unknown structural information and hidden evolutionary features^7^. In addition, a series of language models such as UTR-LM^10^, CaLM^11^, CodonBERT^12^ have been applied in the analysis and scoring of mRNA sequences. UTR-LM is trained with 5′ UTR sequences using a BERT-based architecture, incorporating individual nucleotide tokenization within a multi-task learning framework, and achieves the state-of-the-art performance on UTR-related tasks, such as the prediction of translation efficiency (TE)^13^ and mean ribosome loading (MRL)^14^. Notably, CaLM significantly expands the vocabulary of protein language model by substituting the vocabulary of amino acid residues with their degenerate codons, which effectively reduces the entropy of the model and enhances the compression efficiency during text processing.

Based on an architecture of transformer prefix decoder^15^, here we develop a language model of the integrated protein and mRNA coding sequences, termed BiooBang (Fig. 1). The sequences of CDS and encoded protein are treated as natural language and applied to self-supervised learning for training. BiooBang exhibits excellent performances in a couple of discriminative tasks, including TE and MRL predictions compared to the models that only utilize one modality, either amino acid sequences of protein or nucleotide sequences of CDS, as the sole input at the pre-training stage. Moreover, it also possesses the generation capability in an autoregressive manner similar to GPT, which is further validated via the wet-lab experiments in the significantly higher yield of recombinant protein using the BiooBang-generated CDS candidates.

**Fig. 1.**
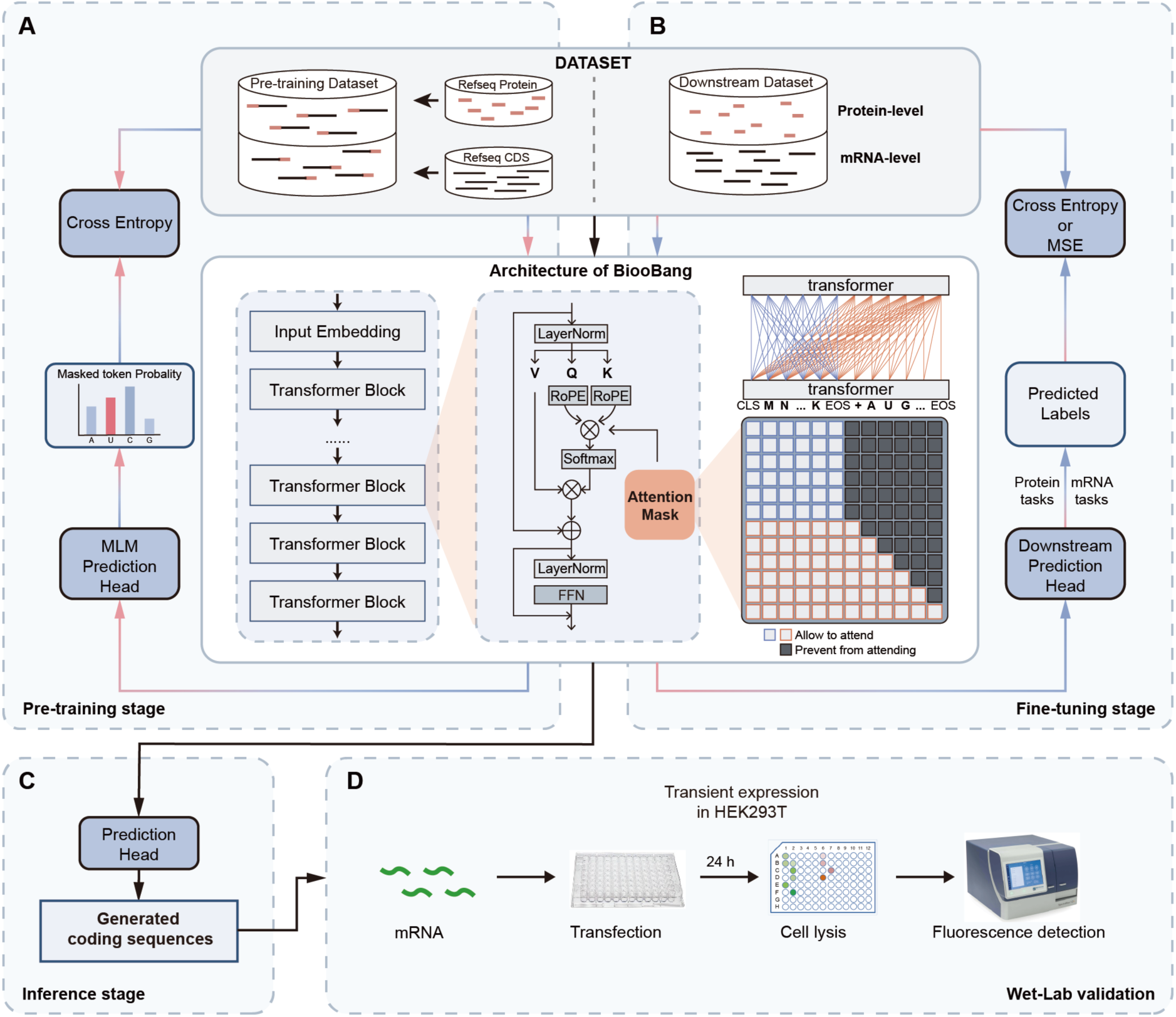
A flowchart of BiooBang model and wet-lab validation. (**A**) At the pre-training stage, the protein and corresponding coding sequences (CDS) or the reverse combinations are tandemly input to the model, enabling the performance of modeling on two types of sequences simultaneously. Via designing an attention mask matrix, the model is equipped with the ability for bidirectional understanding and unidirectional generation. (**B**), At the fine-tuning stage, the model is fine-tuned using either proteins or mRNA sequences according to the corresponding task. (**C**), At the inference stage, a given protein sequence is input to the model, which autoregressively generates dozens of coding sequences (50 sequences in our case) through the prediction head. (**D**), The wet-lab validation of the generated sequences in HEK293T.

In summary, the integration of protein and mRNA coding sequences enables BiooBang not only to mutually enhance semantic compression, but also exhibit the improvement of the capacity in optimizing the coding sequences for the desired purpose. Our findings provide profound insights into the yet-unknown intrinsic correlations between protein and mRNA sequences as well as between CDS and 5’ UTR sequences, implicating the promising applications in synthetic biology and biomedicine.

## Results

### Architecture of BiooBang enables simultaneous integration of both protein and CDS sequences

Based on an architecture of transformer-based prefix decoder^15^, here we develop a language model of the integrated protein and mRNA coding sequences, termed BiooBang (Fig. 1). We pretrained the foundation model, using the randomly chimeric protein (in amino acid residues) and coding (in nucleotides) sequence pairs from the RefSeq database^16^. We applied an adjustable attention mask matrix, enabling the model feasible to various tasks upon different processing strategies of input data (Fig. 1). In addition, we utilized not only the bidirectional attention to capture the global context of the sequences, but also the unidirectional attention to ensure the consistency of autoregressive generation contexts. Simultaneous input of paired protein-CDS sequences ensures the strict adherence to the rules of codon translation between the tokens of amino acid residues and nucleotides, making the model capable of capturing the syntactic patterns in both protein and coding sequences.

Different from the popular PLMs or RNA/codon language models, we model two modalities, protein and CDS sequences, in a combinatory manner using shared parameters and architecture. The shared parameters might make the model gain the capability of learning more generalized representations that are jointly optimized with various tasks, allowing to decipher the intrinsic disciplines between two modalities. Notably, the fusion of bidirectional and unidirectional attention architectures in context is able to effectively mitigate the risk of overfitting which often occurs in unimodal models. Moreover, compared to most previous biological language models, BiooBang possesses the capability on both discriminative and generative tasks.

### Performance of the model in predicting the physiochemical parameters of protein and mRNA

Based on the foundation model, we employed a couple of standard downstream tasks to test the performance of the model (Fig. 1B). These tasks are six protein-related^17–27^ and four mRNA-related^28–31^, respectively. The six protein-related tasks include the prediction of GFP fluorescence (FluP)^17,18^, GB1 fitness (GB1P)^19^, protein solubility (SolP)^20^, protein folding (FoldP)^21^, secondary structure (SSP)^22–26^ and subcellular localization (LocP)^27^. As we know, FoldP and SSP are used to assess the semantically capture of the protein structure information, with FoldP referring to multiclass classification on the folding structure, and SSP involved in the multilabel classification at the token-level. Notably, the model was trained using a frozen-parameter fine-tuning to validate the representational capacity of the foundation model (Fig. 2A). We found that BiooBang exhibits a better performance on tasks GB1P, FluP, and FoldP, compared to the previous benchmark models such as ESM2^7^, ProtT5-XL-U50^8^ and Ankh^9^ (Fig. 2B and table S1). BiooBang achieves a Spearman’s *ρ* of 0.905 for GB1P and 0.662 for FluP, and an accuracy of 69.1% for FoldP. These values are even higher than those of the best-performing model Ankh, which possesses 1.15 billion parameters. Concerning SSP task, BiooBang shows a comparable performance to all PLMs, except for a slightly lower Spearman’s *ρ* compared to Ankh.

**Fig. 2.**
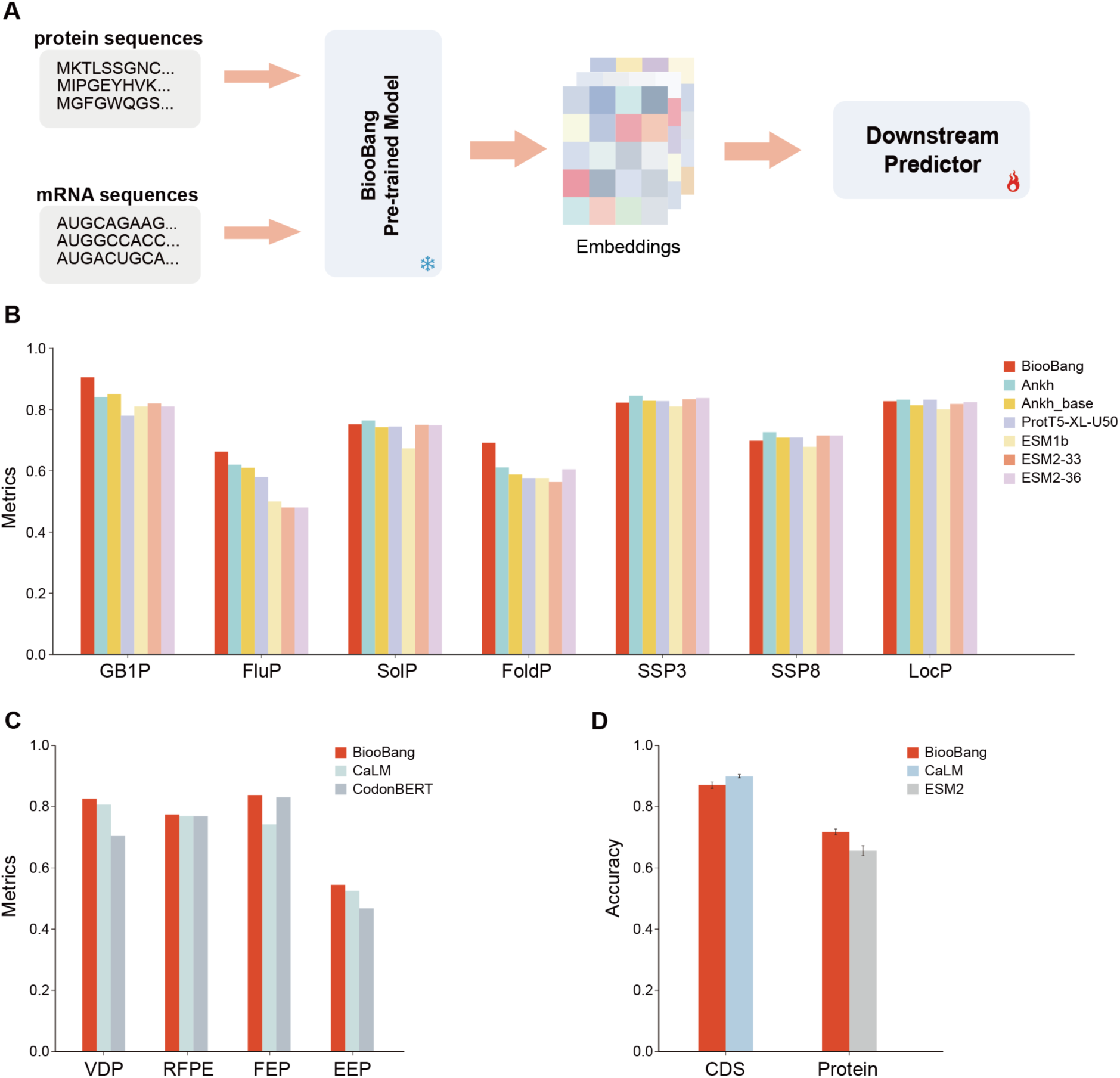
Evaluation the performance of BiooBang on protein and mRNA-related tasks. (**A**) The fine-tuning approach utilizes a frozen-parameter strategy 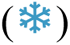, ensuring that the pre-trained model’s parameters remain fixed until the downstream predictors have been well trained 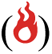. (**B**) The performance of the model on protein-related tasks. The tasks include the prediction of GB1 fitness (GB1P), GFP fluorescence (FluP), protein solubility (SolP), protein folding (FoldP), secondary structure (SSP) and subcellular localization (LocP). GB1P and FluP are regression tasks, evaluated by Spearman’s *ρ*; whereas the others are classification tasks, evaluated by the accuracy. (**C)** The performance of the model on mRNA-related tasks. SARS-CoV-2 Vaccine Degradation Prediction (VDP), mRFP Expression Prediction (RFPE), and Fungal Expression Prediction (FEP) are regression tasks, evaluated by Spearman’s *ρ*; whereas Escherichia coli Expression Prediction (EEP) is a classification task, evaluated by the accuracy. (**D**) The accuracy of nearest-cluster-centre classification. The inputs of CDS and protein sequences are evaluated, respectively. BiooBang has the capacity to accept two distinct types of corpora. In contrast, CaLM and ESM2 focus only on one type of corpora, either CDS or protein.

For the mRNA-related tasks, we processed data by the partitioning method according to CodonBERT^12^, and compared to the benchmark models^11,12^. The four tasks include the prediction of expression in Escherichia coli (EEP)^28^, as well as mRFP (RFPE)^29^, fungi (FEP)^30^, and SARS-CoV-2 vaccine degradation (VDP)^31^. BiooBang achieved a higher accuracy of 0.545 in EEP and a higher Spearman’s *ρ* of 0.775, 0.838, and 0.827 in the other tasks (Fig. 2C and table S2). Thus, BiooBang has the best performance compared to the two codon language models CaLM and CodonBERT.

We further conducted dimensionality reduction and nearest-cluster-centre classification to evaluate the model, via comparing to the codon and protein language models, respectively (Fig. 2D). With the same input of protein sequences, the classifier of BiooBang yields an accuracy of 71.78% (Fig. 2D and fig. S1), 9.4% higher than that of ESM2^7^; whereas, upon inputting the same coding sequences, BiooBang gains an accuracy of 87.03%, slightly lower than that of 89.96% for CaLM^11^. It indicated that upon incorporation of CDS at the pre-training stage makes our model gain the capacity to decipher more hidden intrinsic information in the protein sequences. Furthermore, the integration of protein and coding sequences significantly elevates the adaptability and performance of our model on understanding both protein and CDS contexts.

### Application of fine-tuned models to the prediction of TE and MRL

Elevated translation efficiency of mRNA is indispensable for the rapid and precise production of the desired proteins, especially for those commercialized proteins in the bioengineering and biopharmaceutical fields. Recent omics techniques enable us to quantitatively determine the TE of genome-level mRNA molecules, described as the ratio of the normalized reads (reads per kilobase per million-mapped reads, RPKM) of ribosomal footprints on the mRNA (Ribo-seq) over RNA sequencing (RNA-seq)^13^. The experimental data of Cao et al.^13^ display a long-tailed distribution of TE values in the three specific cell lines (fig. S2), suggesting that most mRNA data exhibits moderate or low translation efficiency, with a few of high efficiency. Once applying the TE values to a logarithmic transformation^10,13^, the resulted LnTE data will become more statistically valid for the subsequent model building and analysis.

The foundation model was fine-tuned on three endogenous datasets of human muscle tissue (Muscle), human prostate cancer cell line PC3 (PC3), and human embryonic kidney (HEK) 293T cell line^13^, using the sequences of CDS and 5′ UTR as fine-tuning inputs to produce the BiooBang-CDS and BiooBang-UTR models, respectively. For the downstream predictor, we applied a multilayer perceptron (MLP) with an architecture the same to UTR-LM^10^. In both CDS and 5′ UTR inputs, BiooBang achieves better performance with all three datasets on TE prediction task compared to UTR-LM (Fig. 3, A and B and table S3). In details, BiooBang-CDS outperforms UTR-LM, which is pre-trained on 5′ UTR sequences, by ∼37% in Spearman’s *ρ*. Moreover, out of our expectation, the foundation model of BiooBang, which was solely pre-trained on CDS, also exceeds UTR-LM by ∼13% upon being fine-tuned with 5′ UTR data.

**Fig. 3.**
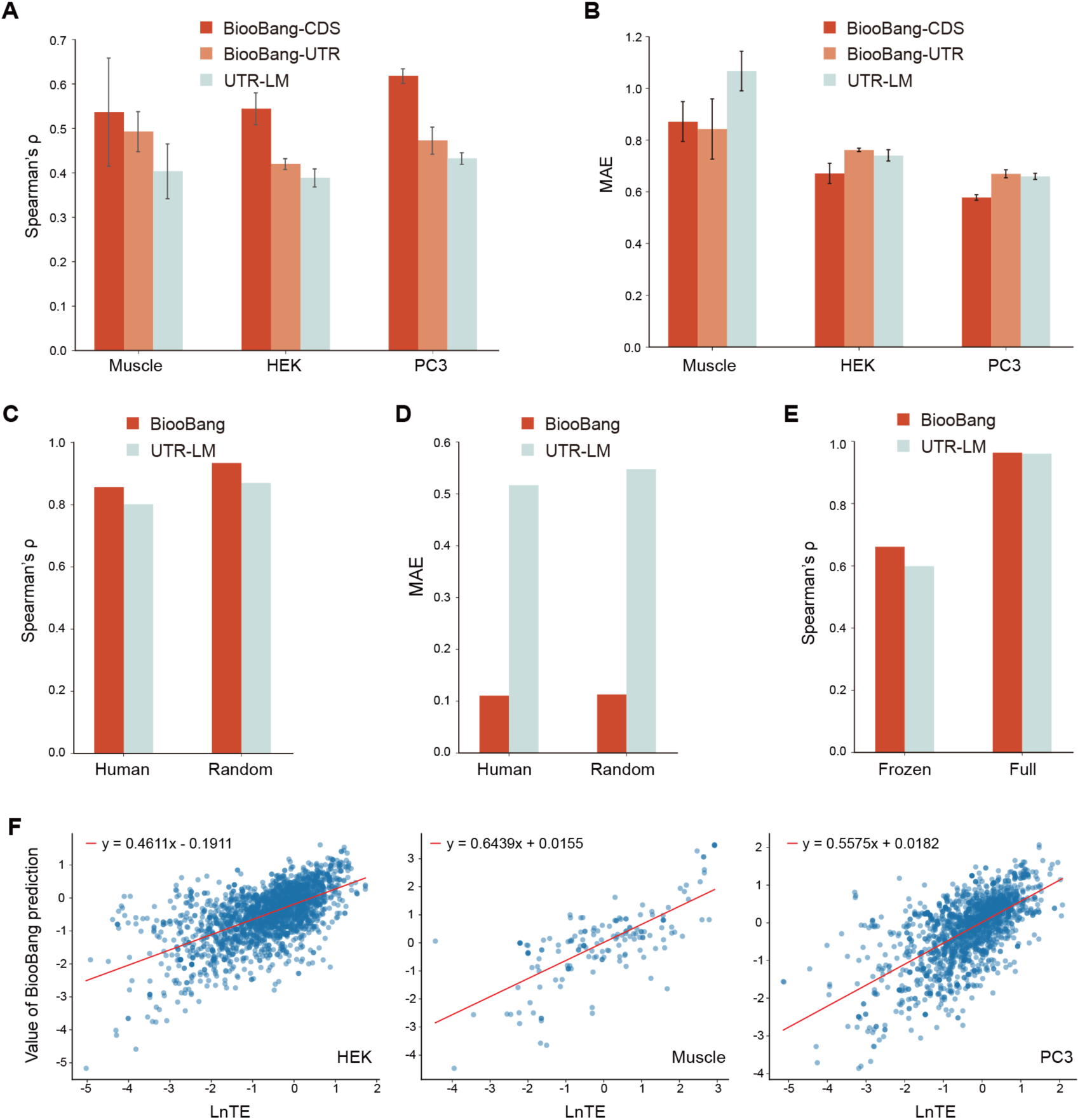
Evaluation of prediction on the translation efficiency (TE) and mean ribosome loading (MRL). (**A** and **B**) The TE prediction evaluated by (A) the Spearman’s *ρ* and (B) the mean absolute errors (MAE), in various cell lines (muscle, HEK293T and PC3). BiooBang-CDS and BiooBang-UTR are fine-tuned using the coding sequences (CDS) and 5′ UTR sequences, respectively. Data are presented as mean values ± s.d. obtained via fivefold cross-validation. (**C** and **D**) The MRL prediction evaluated by (C) the Spearman’s *ρ* and (D) the MAE. (**E**) Ablation study of two fine-tuning strategies with frozen or full parameter. (**F**) The prediction of BiooBang-CDS on TE (normalized in LnTE) in HEK293T, Muscle and PC3 cell lines. In these cell lines, 2315, 160 and 1800 points in the validation set are presented respectively. The linear regression equations of the scatter plot are shown as the inlets. Please do not use text boxes to arrange figures. High-resolution (preferably editable PDF or Adobe Illustrator format) figure files will be requested following review.

As MRL is an indicator of translation efficiency for specific mRNA that is primarily influenced by the 5′ UTR, in consequence we investigated the representation capability of BiooBang in 5′ UTR sequences on the MRL task^14^. Different from that of the TE datasets, the MRL datasets were proceeded using variable 5′ UTR sequences and a fixed CDS. By employing the independent tests on a dataset containing 83,919 random 5′ UTRs and 15,555 human 5′ UTRs, BiooBang demonstrates significant improvement over UTR-LM, with a 7% increase in Spearman’s *ρ* and a reduced MAE to ∼1/5 (Fig. 3, C and D and table S4). Moreover, we designed an ablation study on the training strategy with U1 library^14^, showing that our model has an increased Spearman’s *ρ* of ∼10% compared to UTR-LM using frozen parameter fine-tuning (Fig. 3E and table S5). The results indicate that there exist not only some obvious correlations between CDS and TE, but also yet-unknown crosstalk between 5′ UTR and succeeding coding sequence.

In fact, our model essentially serves as a function indicator for artificial fitting, thus we also compared our model with the traditional strategies for protein expression optimization, including codon adaptation index (CAI)^32^, minimum free energy (MFE)^2,33^ and MFECAI^34^. CAI and MFE are based on the codon usage preferences in a given species and RNA secondary structure, respectively; whereas MFECAI is a joint optimization objective of LinearDesign that combines CAI with MFE by employing lattice parsing. Compared to aforementioned strategies, the prediction results of BiooBang-CDS exhibits a much more significant correlation to LnTE (Fig. 3F and fig. S3). One explanation is that the traditional strategies only consider the codon usage preferences at the species level, without counting the tissue specificity^35^.

### De novo generated coding sequences of high TE and experimental validation

Nowadays, optimization of protein expression is only a compromise solution based on the heuristic rules without fully decoupling gene information. In contrast to the traditional strategies, our methodological framework is independent of any predefined disciplines of codon usage preferences or mRNA stability. Our model possesses a prefix-decoder architecture that endows it with the generative capability, which could be applied to de novo generation of coding sequences based on comprehension of the encoded protein sequences from the latent space.

In the past decades, transcriptomics investigations on various tissues indicated that the existence of diverse and specific gene expression patterns^36^, which was proposed to possess some yet-unknown high-dimensional syntactic features of tissue specificity. To produce desired coding sequences based on the generative capability of the model, the RNA sequences of high TE could be input to the foundation model for the supervised fine-tuning (SFT) (Fig. 4A). Here we collected hundreds of mRNA sequences of high TE score expressed in HEK293T from the transcriptomics data^13^, to construct the fine-tuned model of generative capacity. Notably, both the masked language modeling (MLM)^5^ and causal language modeling (CLM)^6^ were applied during the fine-tuning stage (Fig. 4B). In both tasks, we utilized the same strategy for self-attention mask matrix as GPT^6^, allowing the model to focus solely on the preceding context for predicting the next token.

**Fig. 4.**
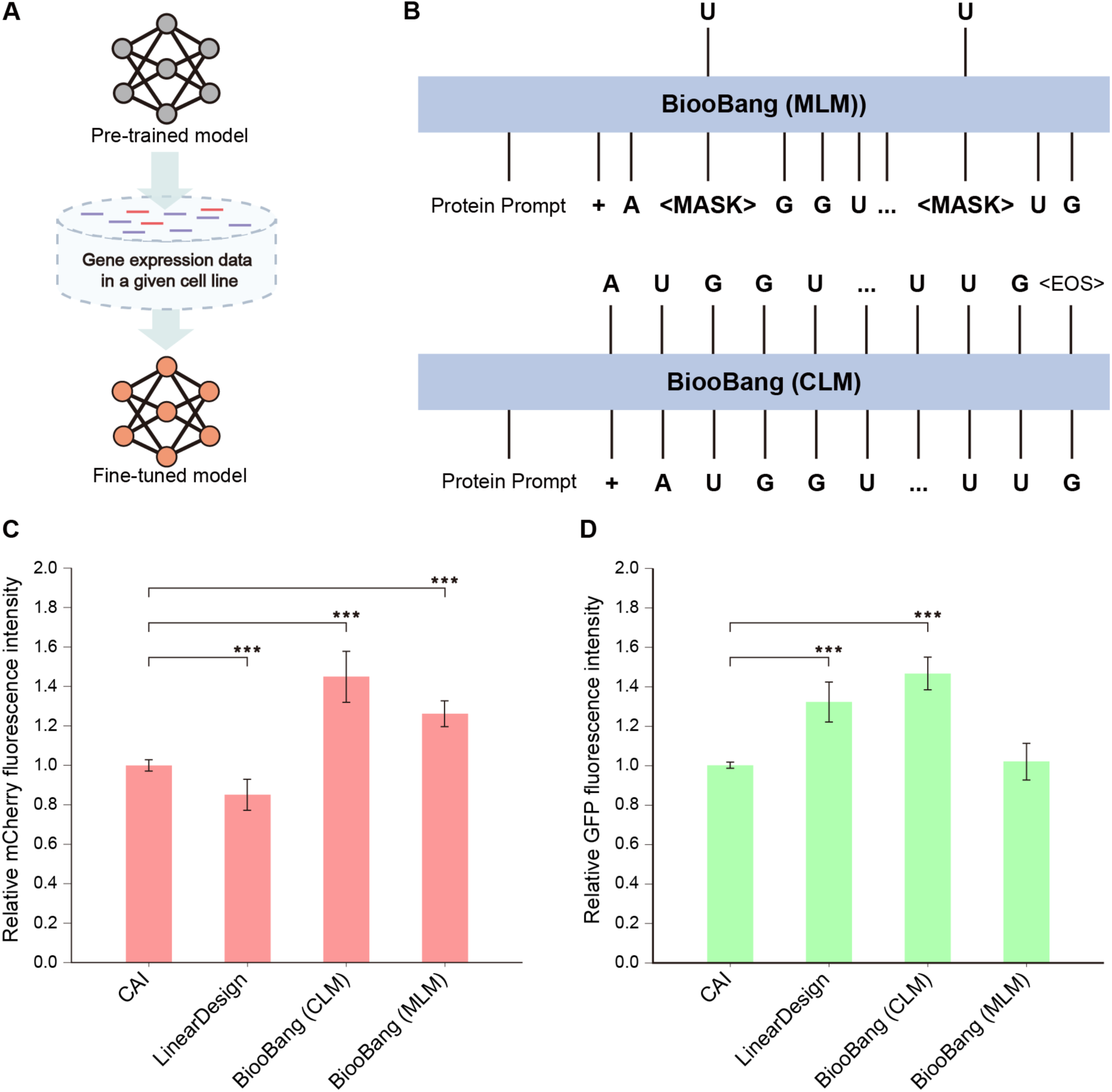
The fine-tuned generative model and wet-lab validations. (**A**) The gene expression data in a given cell line were used for producing the fine-tuned model. In the present case, the high TE dataset from HEK293T were used. (**B**) The schemes of the masked and causal language modeling (MLM and CLM). The MLM predicts masked tokens, whereas the CLM predicts the next token using a shifted prediction approach without masking the data. (**C** and **D**) The relative fluorescence intensity of HEK293T cells following transfection with different designed mCherry (C) or GFP (C) mRNA. The output intensity of traditional strategy (CAI) was set to 1.0. Three independent assays were performed to calculate the means and standard deviations, and the data are presented as the means ± S.D. The two-tailed t-test was used for the comparison of statistical significance. The p values of < 0.001 were indicated with***.

To test the autoregressive generation capability, we selected the commonly used reporter genes mCherry and GFP as the examples. Following the traditional strategy, the coding sequences of mCherry and GFP were usually designed via setting a CAI of 1.0, which means the single codon of highest frequency was assigned to a given amino acid. The sequences optimized by LinearDesign, upon setting a hyperparameter *λ* of MFECAI at 0.5, also serve as the benchmark. Using the beam search, BiooBang (CLM and MLM) generated a series of sequences, one of which possesses the highest total probability of tokens was applied to experimental validation.

In total, four coding sequences were designed via various strategies, synthesized and in vitro transcribed into mRNA, which were subsequently translated in HEK293T (Fig. 1D, fig. S4 and S5). Compared to the traditional strategy based on CAI, the sequences designed by LinearDesign exhibit a lower expression of mCherry (∼15% decrease) and a higher expression of GFP (∼32% increase) (Fig. 4, C and D). In contrast to the traditionally designed sequences of 1.0 CAI, the generated sequences of BiooBang via causal language modeling show a ∼45% elevated expression level of both mCherry and GFP. However, the sequences of BiooBang via masked language modeling yield a better performance (∼25% increase) in mCherry, but comparable in GFP expression. Taking the benchmark sequences of LinearDesign as the controls, the coding sequences generated by BiooBang (CLM) display an increase of ∼60% and 10%, in mCherry and GFP, respectively. The difference between CLM and MLM of BiooBang might be attributed primarily to the higher training efficiency of CLM, each token of which has been iteratively trained; whereas only the masked tokens have been trained in MLM.

## Discussion

Via a biological foundation model that unified both discriminative and generative tasks, for the first time we explicitly integrate the synergistic information of protein and mRNA coding sequences through dual modeling. It enables our model to efficiently address diverse downstream tasks, no need to modify the architecture of either protein or CDS contexts. Notably, compared to the protein-only language models, our model exhibits a significantly better profile especially on the prediction tasks of GB1P and FluP, which focus on predicting the activity of protein variants using the random mutation strategy. It indicates that incorporation of CDS information at the pretraining stage distinctly promotes the capability of our model to understand the primary sequences of proteins at the fine-tuning stage. This unexpected finding is potentially rooted from the shared token-level rules between protein and coding sequences, which gains the supremacy over the synteny of CDS and protein based on the Central Dogma and codon degeneracy. It is conceivable that the as-yet unknown high-dimensional information embedded in mRNA sequences is intrinsically connected to the biological features of protein sequences.

In addition, the prediction of TE or MRL using 5′ UTR sequences shows a better profile than the state-of-the-art model (UTR-LM), indicating that our model possesses the ability to generalize from the CDS to its upstream 5′ UTR. Therefore, we hypothesize that there should exist some intrinsic crosstalk, but yet-unknown, between the non-coding and coding regions, which have been preserved along the long history of co-evolution. Hopefully, it might provide a new paradigm that leverages the transferability and generalization capability of the pre-trained model to interpret the syntactic and semantic correlations between the two distinct corpora, which could be further applied to mine more unknown correlations among other forms of biological information.

Upon learning the intricate relationships between various sequence elements, our model also gains the ability to translate these knowledges into de novo generation of desired sequences. Moreover, two examples of generated coding sequences were validated by the positive feedbacks of wet-lab experiments. Thus, one potential direction is the bona fide reinforcement learning from human feedback (RLHF)^37,38^, which could iteratively align the sequences generated by the model with those the human desires.

In conclusion, the performance of our model on various biological tasks reveals the potential to integrate various features or biological “languages” in the foundation model. The crosstalk between these distinct sequence modalities might enable us to uncover the hidden and interdependent layers of complex biological systems. It will not only advance the prediction capability, but also unravel at the higher dimension the intricate relationships that have been long overlooked by the present single-modality approaches. The synergy between various biological “languages” may represent a frontier for advancing computational biology in an unprecedented way.

## Methods

### Data

#### Evaluation Dataset for Species identification

For the dimensionality reduction and visualization, we used the species dataset reported by CaLM^11^, which includes sequences from seven model organisms across the tree of life (fig. S1): three multicellular eukaryotes (*Arabidopsis thaliana*, *Drosophila melanogaster*, and *Homo sapiens*), two unicellular eukaryotes (*Saccharomyces cerevisiae* and *Pichia pastoris*), bacteria (*Escherichia coli*), and archaea (*Haloferax volcanii*).

For the nearest-cluster-centre classification, we utilized another species dataset described in CaLM, consisting of 4,358 sequences. After removing long sequences, the dataset was reduced to 4,032 sequences. We then randomly divided the dataset into five subsets for 5-fold cross-validation.

#### Downstream datasets for protein-and mRNA-related predictions

We validated the performance of the model in predicting the physicochemical properties of proteins and mRNA across six protein-related tasks and four mRNA-related tasks.

For the protein-related tasks, we followed the task selection outlined in Ankh^9^, which includes Fluorescence Prediction (FluP), Solubility Prediction (SolP), GB1 Fitness Prediction (GB1P), Fold Prediction (FoldP), Secondary Structure Prediction (SSP), and Sub-cellular Localization Prediction (LocP). Moreover, we selected four mRNA related tasks, *E. coli* Expression Prediction (EEP), mRFP Expression Prediction (RFPE), Fungal Expression Prediction (FEP) and SARS-CoV-2 Vaccine Degradation Prediction (VDP), as described in CodonBERT^12^.

In details, FluP utilized the fluorescence intensity dataset of green fluorescent protein mutants annotated by Sarkisyan et al.^17^, following the dataset splits used in TAPE^18^. GB1P was used for the regression of GB1 binding fitness after mutations at four specific positions, using the dataset and splits from FLIP^19^. SolP employed the solubility dataset used in the development of DeepSol^20^, adhering to the same data splits. FoldP classified proteins into one of 1,195 possible folds, utilizing the dataset and original splits provided by Hou et al.^21^. SSP is a token-level task divided into Q3 and Q8, classifying amino acid residues into three or eight-fold classes, respectively.

The training set of SSP was derived from NetSurfP-2.0^26^, with test sets including CASP12^23^, CASP14^24^, TS115^25^, CB513^22^. LocP utilized the DeepLoc^27^ dataset, following the same data splits. EEP was derived from the MPEPE dataset^28^ and used to predict protein yields in *E. coli*. RFPE was based on the dataset from Nieuwkoop et al.^29^, predicting the expression of several gene variants in *E. coli*. FEP utilized the dataset provided by Wint et al.^30^, which included a wide range of fungal CDS and tRNA sequences. VDP was sourced from the dataset by Leppek et al.^31^, combining structural features, mRNA stability and their translation efficiency to predict degradation in SARS-CoV-2 vaccines. The data splits for all mRNA-related tasks were consistent with those used in CodonBERT, ensuring comparability in different models.

For the TE prediction, we utilized the dataset from Cao et al.^13^, which includes three endogenous datasets: HEK, Muscle, and PC3. We downloaded the corresponding coding sequences based on the provided ENST IDs. The sequences were further processed by removing any coding sequences with multiple stop codons or those whose lengths were not multiples of three. To prevent information leakage, we used MMseqs2 tool^39^ to cluster the sequences at 50% identity, with the specific parameters “--cov-mode 0 -c 0.8 –min-seq-id 0.5 -s 7.5”. We then divided the clusters into 5 folds for cross-validation. For the MRL prediction, we used the dataset from Paul et al.^14^ and conducted ablation experiments on the U1 library, which has been widely used in UTR researches. The row data distribution of TE and MRL exhibited a long-tail distribution, thus we applied logarithmic normalization to approximate a near-normal distribution (fig. S2).

In addition, we performed independent tests on two 5’ UTR datasets: one containing 83,919 random 5’ UTR sequences and another with 15,555 human 5’ UTR sequences. We followed the data split used by Optimus^14^, where the test set included 7,600 random 5’ UTRs and 7,600 human 5’ UTRs.

#### Fine-tuning dataset for the generative model

We selected sequences with LnTE > 0.5 from the HEK293T TE dataset^13^ to fine-tune the generative model for coding sequence. Meanwhile, we clustered these filtered sequences using MMseqs2^39^ to prevent overfitting during the training period. We randomly selected 20% of the clustered sequences as the validation set, while the remaining 2,560 sequences were designated for the training set.

### Model architecture

#### The architecture of BiooBang

BiooBang is a transformer-based biological language model primarily trained on protein and mRNA coding sequences (Fig. 1). Similar to other language models, the core architecture of BiooBang consists of two main components: an embedding module and transformer blocks^4^. Our tokenizer converts input string sequences into token IDs. For protein sequences, we have inherited the vocabulary from ESM2. Besides protein tokens, we have added five additional characters—A, U, C, G, and N—to the vocabulary for mRNA coding sequences. Regarding special tokens, “CLS” is used as the start token, “<EOS>” as the end token, “<MASK>” as the masking token, “+” as the generation prompt token, and “<PAD>” as the padding token.

The embedding module maps input tokens into representation vectors, which are subsequently processed by the transformer blocks. During this process, we employed rotary position embedding (RoPE) integrate positional information into the attention mechanism^40^. Each transformer block comprises an attention mechanism and a feedforward neural network. Consistent with current mainstream models, we employ post-layer normalization (post-norm)^41^ in the transformer blocks (Fig. 1). To align with the pre-training tasks, the top layer of BiooBang includes a simple prediction head from RoBERTa^42^, which functions to transform the output features of the model into a vector matching the size of the vocabulary, thereby enabling the prediction of the probability distribution for each token.

The prefix decoder unifies both discriminative and generative tasks by automatically adjusting the attention mask which controls the relationships between tokens. Specifically, the attention module receives three feature vectors as input: Q (query), K (key), and V (value). The main operations performed are:

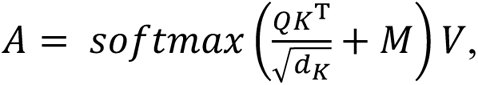

let *M* represent the attention mask. Compared to previous reported biological language models, such as ESM^7^ or ProGen^43^, we concatenated two types of matrices, an all-zero matrix and a strictly lower triangular matrix with negative infinity values in the upper triangle, to define M following the method described in UniLM^15^. This modification not only unifies the training tasks but also significantly improves training efficiency, allowing us to model both bidirectional understanding and unidirectional generation simultaneously using only the masked language modeling (MLM) task.

#### Downstream predictor

We employed three types of predictors for downstream tasks.

For predicting the physicochemical properties of proteins and mRNA, we employed ConvBERT layers^44^ as downstream prediction heads. The parameter configuration of the predictors is identical to that of Ankh, ensuring consistency and comparability in the results.

For the prediction of translation efficiency (TE) and mean ribosome loading (MRL), we adopted the same multilayer perceptron (MLP) used in UTR-LM^10^ as the downstream prediction head. The first linear layer of the MLP is half the size of the foundation model’s output dimension, while the second linear layer corresponds to the number of labels.

For the autoregressive generation of coding sequences with high TE, the prediction head is structurally identical to that used in the pre-training stage, which ensures consistency in functional design, parameter settings, and computational logic between the pretraining and generation stages. This consistency enables the effective use of knowledge from pretraining, leading to more accurate and reliable generation results.

### Training details

#### Pre-training stage

In each training step, the model randomly selects a cluster from the dataset and then chooses a protein sequence (seq-a) along with its corresponding coding sequence (seq-n). To ensure that the inputs adhere to the acceptable length of the model, we implemented a sliding window strategy to truncate seq-a and seq-n while maintaining codon correspondence. The truncated sequences are then concatenated to create the input in the format “<CLS> seq1 <EOS> + seq2 <EOS>” with seq-a and seq-n being placed randomly within seq1 and seq2. Leveraging the prefix-decoder architecture^15^, we unified the pretraining tasks of discriminative models and generative models into the masked language modeling (MLM). For the MLM task, 20% of the tokens were randomly masked, which 80% of the time were replaced with “<MASK>”, 10% with random tokens, and leaving 10% unchanged. The prediction head of BiooBang is then employed to predict the original tokens for each masked position. The objective function for this training is designed to optimize the restoration of these masked tokens, for the seq1:

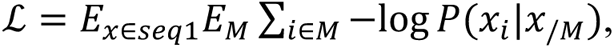

*x* is a single sequence within seq1. M denotes the set of positions where the tokens are masked, and *x_i_* represents the true token at position *i*. The probability *P*(*x_i_*|*x*_/*M*_) corresponds to the prediction of the masked token *x_i_* using all tokens from unmasked positions *x*_/*M*_; for the seq2:

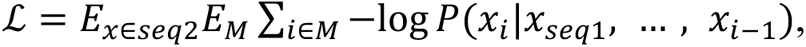

*x* is a single sequence within seq2. M denotes the set of positions where the tokens are masked, and *x_i_*, represents the true token at position *i*. The probability *P*(*x_i_*|*x*_/*seq*1_, …, *x_i_*_-1_) corresponds to the prediction of the masked token *x_i_*, by utilizing the tokens preceding *i* from unmasked positions.

#### Fine-tuning stage

For predicting the physicochemical properties of proteins and mRNA, the model employed ConvBERT layer^44^ as the prediction head and was fine-tuned with frozen parameters. Generally, we utilized the foundation model to generate the embeddings of the target protein or mRNA sequences, and then trained the downstream predictor using these embeddings as inputs.

Following the UTR-LM^10^, we conducted full parameter fine-tuning for the prediction of TE and MRL. To ensure the consistency in the comparative experiments, we utilized the code provided by Chu et al.^10^ to fine-tune UTR-LM on the dataset mentioned above. The final results were obtained through five-fold cross-validation. Especially, frozen parameter fine-tuning was implemented during the fine-tuning stage for the ablation experiments on MRL training strategies. To generate coding sequences with high TE from a given protein sequence, we fine-tuned the model with two strategies: masked language modeling (MLM) and causal language modeling (CLM). The data collator employed in the fine-tuning MLM task was the same as that at the pre-training stage. In contrast, no masking was applied in the fine-tuning CLM task, where the input tokens were shifted one position to the left to serve as the prediction target. Specifically, the optimization function for CLM was defined as:

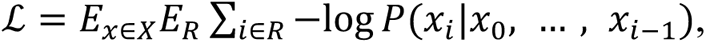

*X* represents a batch of input sequences, where *x* is a single sequence within the batch, and *x_i_*, denotes the true token at position *i*. *R* is a set of position expected to responses, which should be set as the position set of the tokens in the CDS.

#### Unsupervised learning verification

We utilized unsupervised learning to investigate sequence representations across different species, aiming to assess the sensitivity of the pre-trained foundation model to the embeddings of mRNA and protein sequences from various species.

The protein and mRNA sequences were used as inputs to the foundation model, generating the representation embeddings which were subsequently reduced in dimensionality. We then utilized the t-SNE algorithm with the default parameters of Scikit-Learn v.1.3.2^45^ for visualization (fig. S1).

In addition, a nearest-centroid algorithm was applied for species prediction to further validate the representation capability of the model (Fig. 2D). We implemented 5-fold cross-validation for testing, ensuring that the number of sequences in the test set was four times that of the parameter estimation set. Centroids were calculated as the median of the embedding vectors for sequences of each species in the parameter estimation set. During the testing phase, sequences were assigned to the species label of the centroid with the minimum L2 distance.

Based on the two methods mentioned above, we also applied the embeddings of ESM2 and CaLM as benchmark (Fig. 2D and fig. S1).

#### The parameter settings for traditional strategies of expression optimization

The codon adaptation index (CAI) is defined as the geometric mean of the codon optimality for each codon within the mRNA sequence *r*:

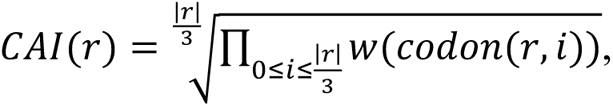

let *r* represent the mRNA coding sequence, and *codon*(*r*, *i*) denotes the *i*-th codon in *r*. In our wet-lab experiments, CAI was employed as one of our baseline strategies to design the coding sequences for mCherry and GFP, with setting it to 1.0. This approach ensured the selection of the highest frequency codon for each corresponding amino acid.

The minimum free energy (MFE) calculation is similarly performed using a structure-based computational method^46^.

MFECAI, an optimization objective used by LinearDesign^34^, is defined as:

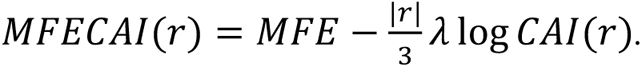

In this research, we set the value of *λ* to 0.5, serving as another benchmark beyond CAI.

### Wet-lab assays

#### In vitro transcription template preparation

All designed GFP and mCherry sequences were synthesized by Sangon Biotech Company and subcloned to the modified vector pUC57 by One Step Cloning Kit (C113-02, Vazyme Biotech co., Ltd). The vector contains a T7 promoter, 5′ UTR, 3′ UTR and a 49-nucleotide poly(A) sequence at the 3′-termini of 3′ UTR. 5′ UTR and 3′ UTR are derived from *human hemoglobin subunit beta*. The synthesized sequences were cloned between the 5′ UTR and 3′ UTR. The linearized vector was obtained with BspQI (10664ES86, Yeasen) restriction and purified by phenol-chloroform extraction followed by ethanol precipitation.

#### In vitro transcription

In vitro transcription (IVT) was performed with the HiScribe T7 high-yield RNA synthesis kit (ON-040, Hongene Biotech) according to the instructions with modifications. In details, a 20 µL transcription reaction contained 1 µg linear DNA template, 8 mM of each NTP, 8 mM of m^7^G(5′) ppp (5′) (2’-OMeA) pG·NH4 RNA cap analog (ON-134, Hongene Biotech), 1.5 µL enzyme mix and 1x reaction buffer. After incubation for 3 h at 37 °C, the DNA was digested by addition of 1 µL DNase I for 30 min at 37 °C. The IVT mRNA was purified using ammonium acetate precipitation followed by ethanol washing. After purification, the mRNA was diluted in 1 mM citrate buffer (AM7001, Thermo Scientific) to the desired concentration and stored at -80℃.

#### Expression assays

HEK293T cells were cultured in Dulbecco’s Modified Eagle’s Medium (DMEM) containing 2 mM L-glutamine, supplemented with 10% fetal bovine serum, 100 U/mL penicillin, and 0.1 mg/mL streptomycin at 37°C in 5% CO2 incubators. Before transfection, HEK293T cells were seeded at 3.0 × 10^4^ cells/well in flat-bottom 96-well plates in 100 µL volume and incubated at 37°C, 5% CO_2_ overnight. 50 ng of GFP/mCherry variants mRNAs were mixed with 50 ng mCherry/GFP mRNA. The addition of a constant amount of mCherry mRNA was used as the reference to avoid any confounding effects during transfection, which allowed us to normalize GFP/mCherry expression to transfection efficiency. The mixed mRNA was transfected using Lipofectamine MessengerMax as per the instructions from the manufacturer (LMRNA008, Thermo Scientific). After 24 h, cells were washed three times using PBS and then incubated with RIPA lysis buffer (R0020, Solarbio) for 15 min at room temperature. The fluorescence intensity was detected on a SpectraMax iD5 (Molecular Devices) plate reader.

## Data and Code availability

The data used for pre-training and fine-tuning the BiooBang model in this study can be accessed on Google Drive (https://drive.google.com/drive/folders/1sQja32qBgx2Z7HD-v_NkT23_RFE2Fs7N) and Zenodo^47^. The pre-training data contains the GCF IDs of all sequences, while the fine-tuning data provides the original sequences and their data splits. The sequences used for experimental validation can be found in https://github.com/lonelycrab888/BiooBang. The BiooBang model will be made available via an open website service for non-commercial use. The supplementary materials provide specific details on the model architecture and training process. Code is not provided.

## Acknowledgments

We thank the developers of PyTorch, MMseqs2, huggingface, biopython, snapgene and others for building invaluable open-source tools and the creators and maintainer of RefSeq, UniProt, Ensembl as well as the researchers whose experimental efforts are included in these resources. Supercomputing Center of University of Science and Technology of China is acknowledged for numerical calculations. This work is supported by the National Natural Science Foundation of China (http://www.nsfc. gov.cn; grant number 32430001), and Anhui Provincial Natural Science Foundation (http://kjt.ah.gov.cn; grant number 202423k09020010). S.-J.H. acknowledges the support from the Postdoctoral Fellowship Program of CPSF under Grant Number GZC20241648. The Supercomputing Center of University of Science and Technology of China is acknowledged for numerical calculations.

## Author contributions

C.-Z.Z., Y.C., S.-J.H. and H.-R.Z. conceived and designed the project. H.L. and E.C. supervised the project. C.-Z.Z., Y.C., S.-J.H., H.-R.Z. and M.-T.C. analyzed data and wrote the manuscript. H.-R.Z. designed and performed machine learning modeling, generation and scoring. J.Z., H.W. and M.-H.F. provided advice on machine learning and computational methods. M.-T.C. carried out the IVT template construction, mRNA preparation, and transient expression assays. B.W., Y.-X.S. and X.-R.Y. performed the overexpression and purification of cytosolic/membrane proteins, and supervised by M.-T.C. All authors discussed the results and assisted during manuscript preparation.

---

**Fig. S1.**
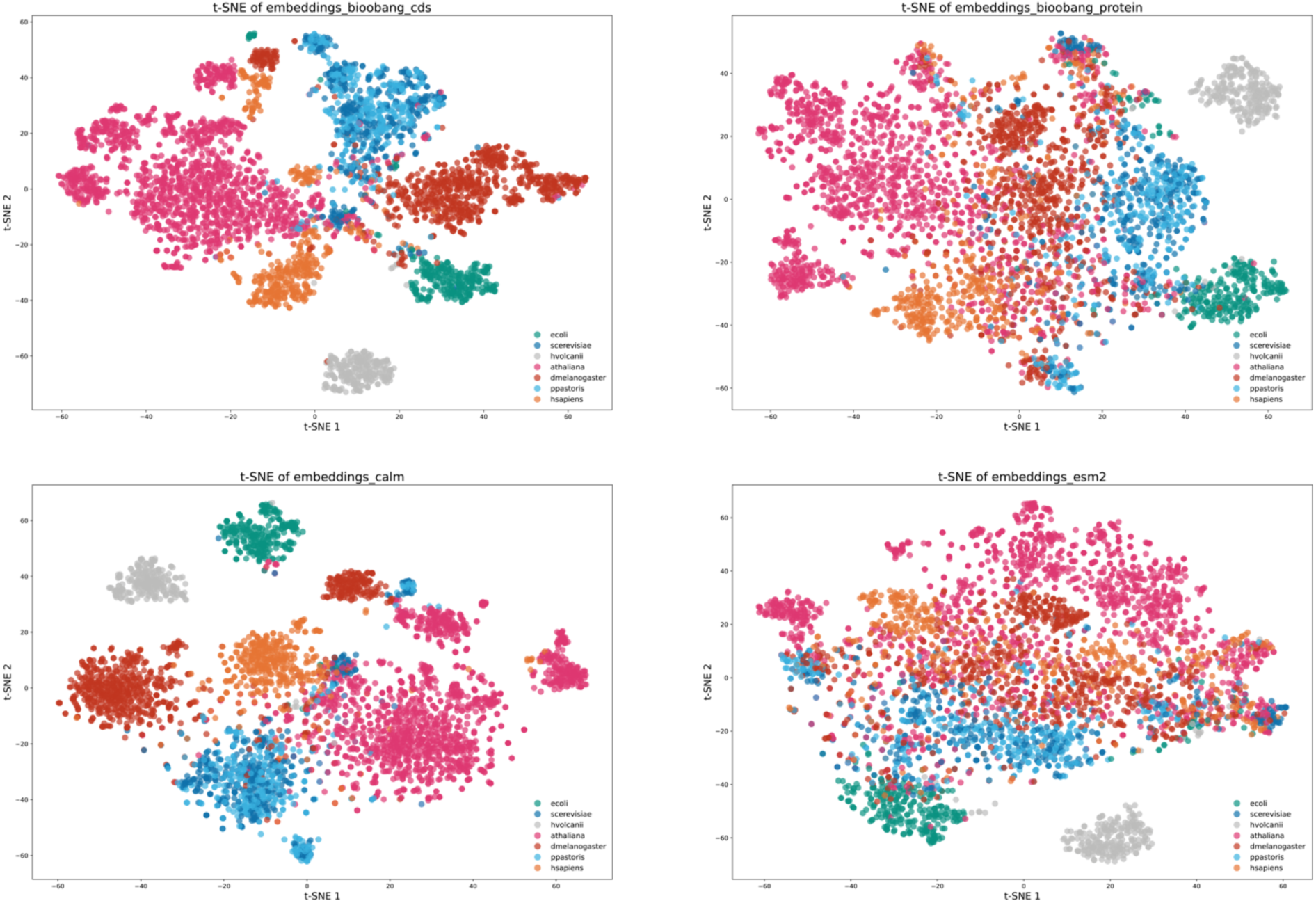
The scatter plot of t-SNE dimensionality reduction in seven species. BiooBang can accept two types of corpora, protein and coding sequences. CaLM and ESM can only accept CDS and protein, respectively. The test species include three multicellular eukaryotes (*Arabidopsis thaliana*, *Drosophila melanogaster*, and *Homo sapiens*), two unicellular eukaryotes (*Saccharomyces cerevisiae* and *Pichia pastoris*), a bacterium (*Escherichia coli*), and an archaeon (*Haloferax volcanii*).

**Fig. S2.**
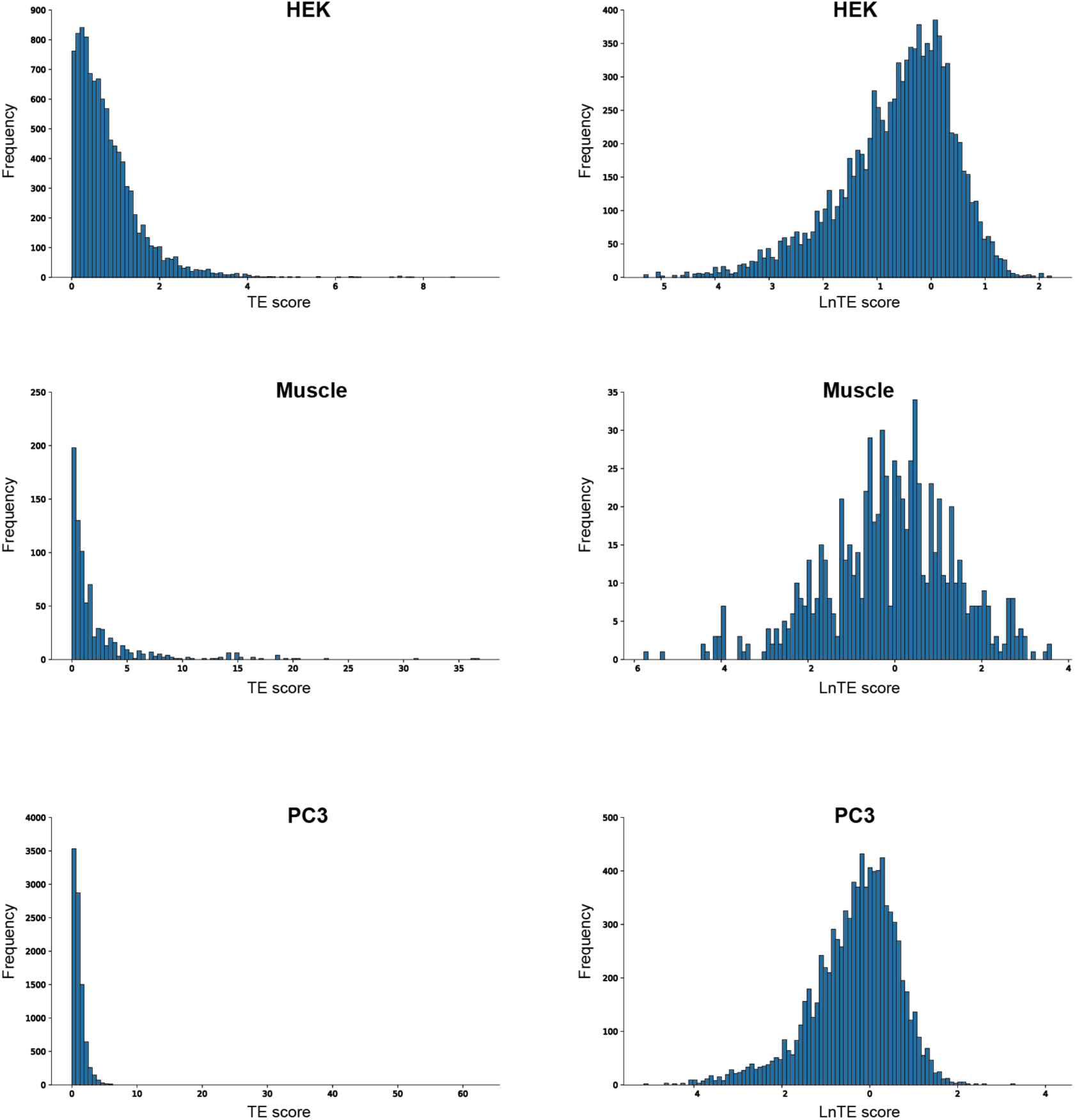
The initial and normalized distributions of TE scores from three cell lines. The raw TE scores exhibit a long-tailed distribution, while the normalized TE scores (LnTE) follow an approximately normal distribution.

**Fig. S3.**
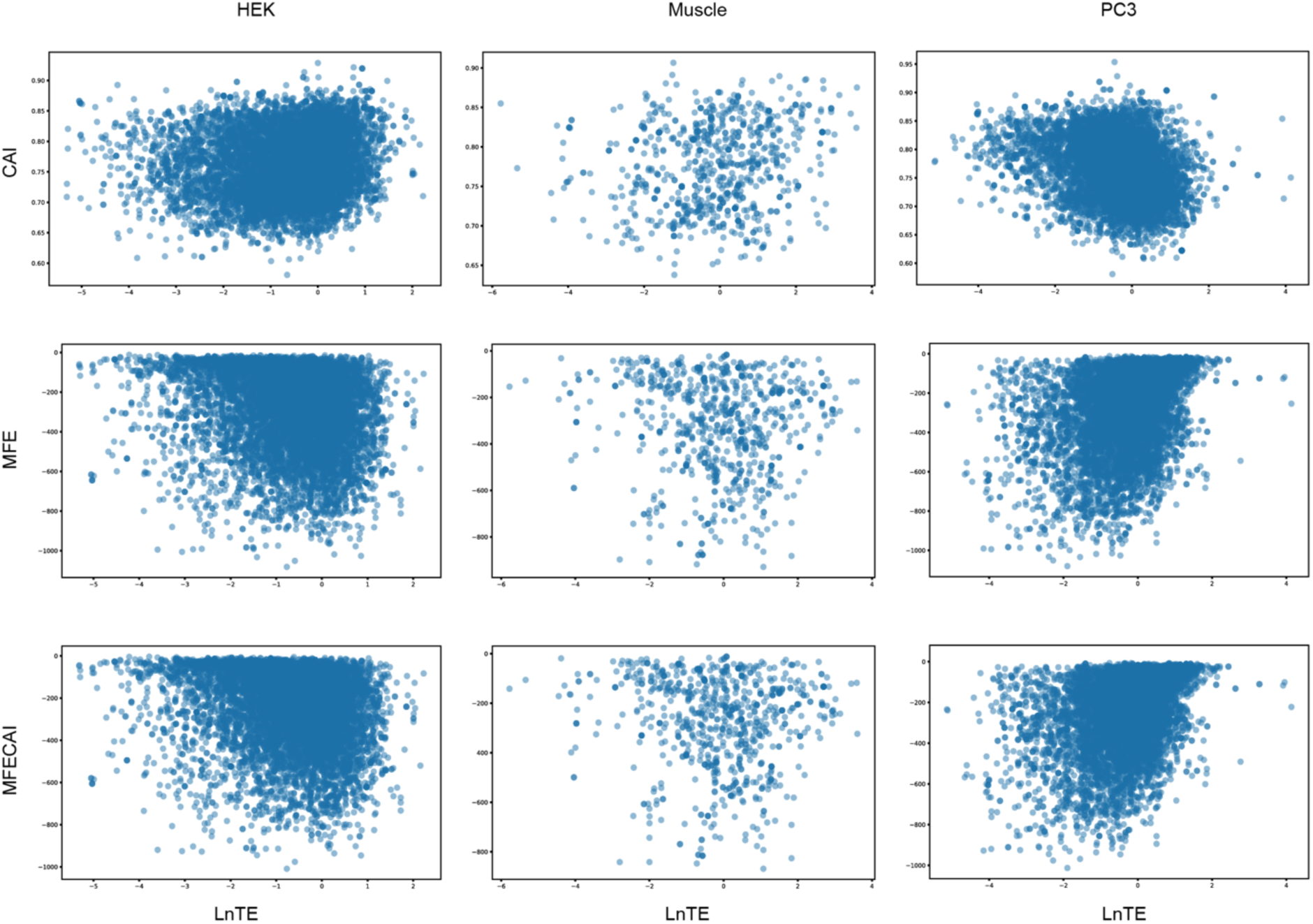
The scatter plot of the scores between LnTE and three popular codon optimization strategies from three cell lines. The objective function of the three popular strategies are CAI (upper), MFE (middle), and MFECAI (lower), respectively.

**Fig. S4.**
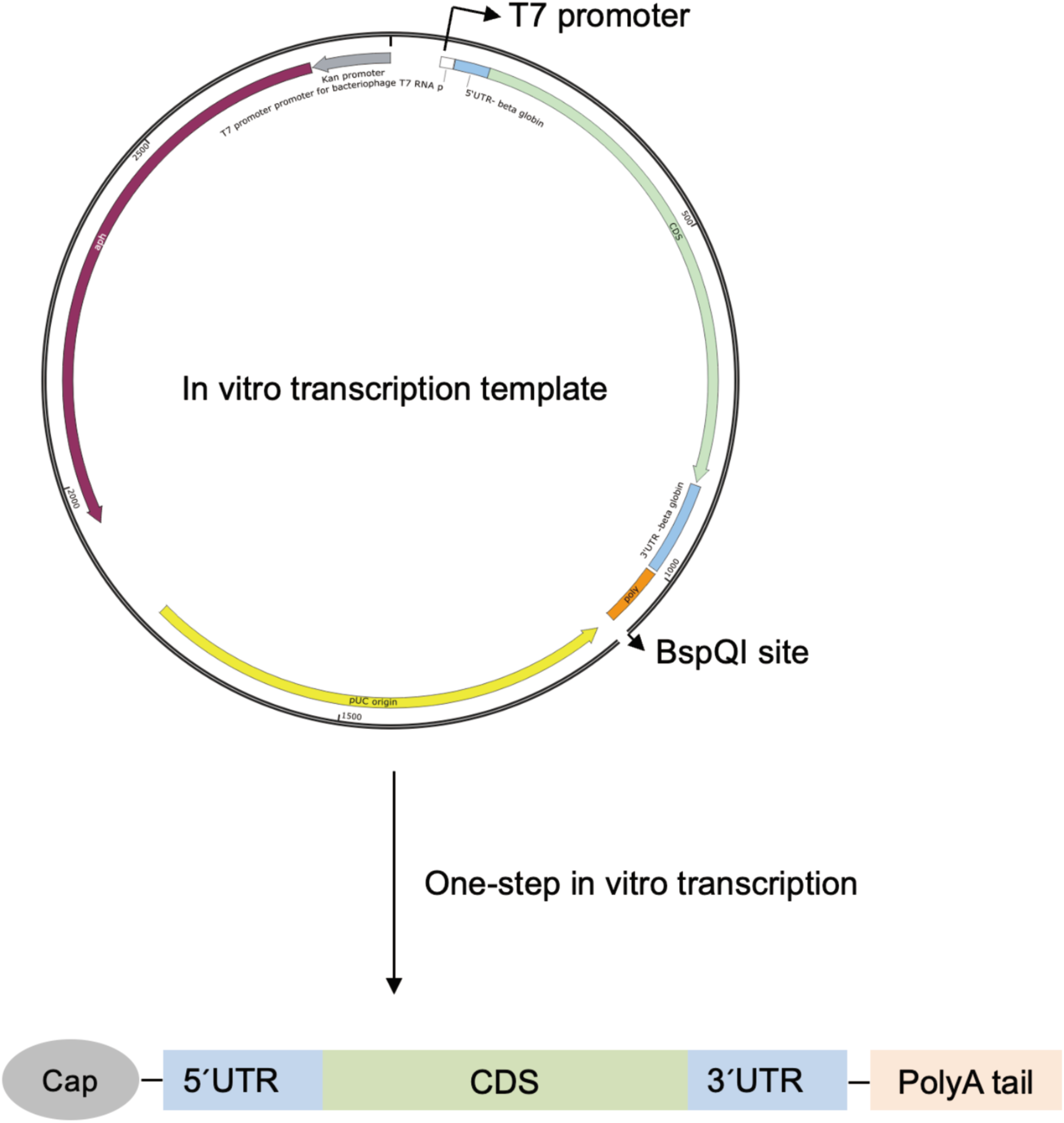
The structure of the in vitro transcription template and the corresponding mRNA.

**Fig. S5.**
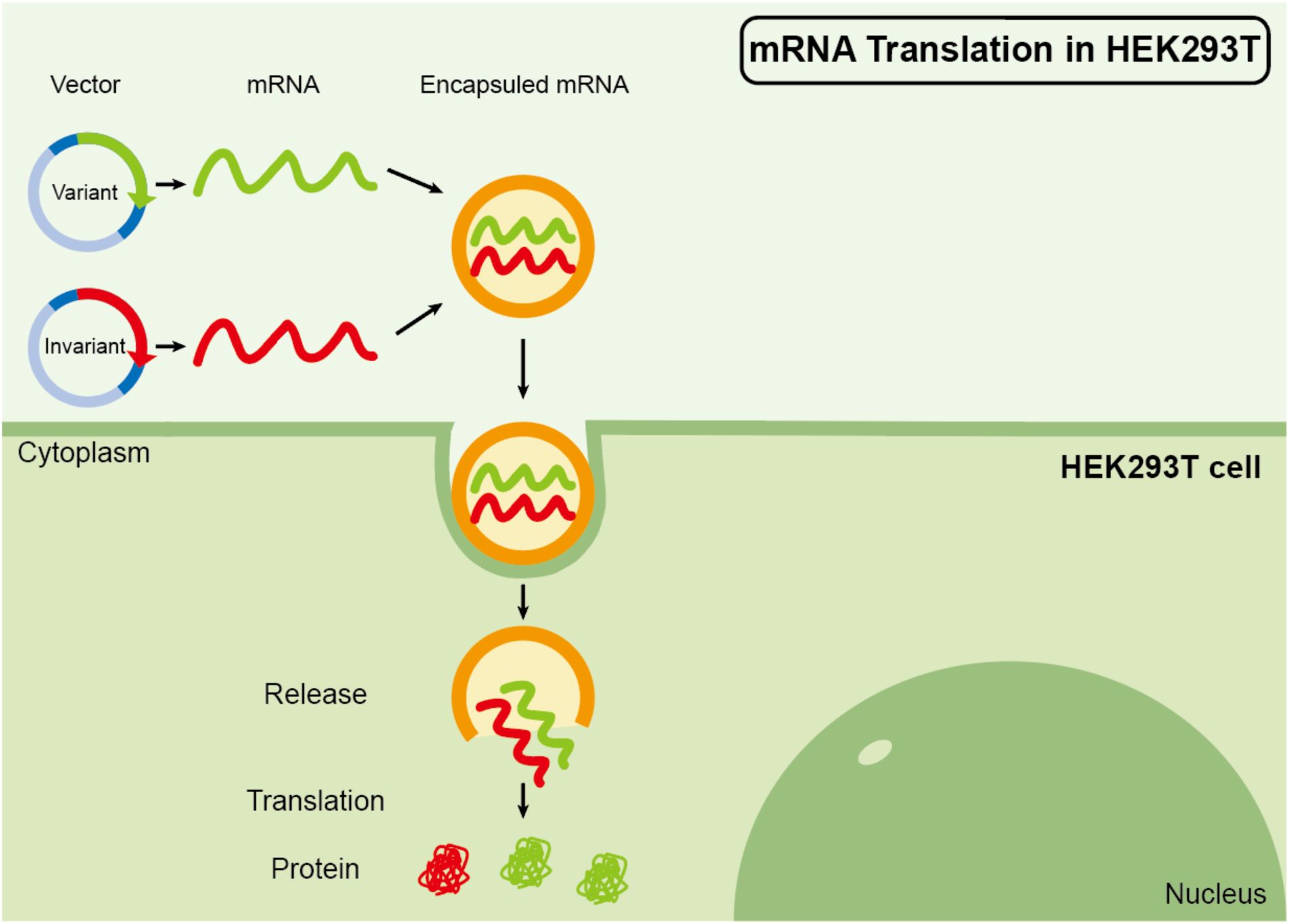
A workflow for mRNA transfection. The mRNA produced by invariant vector was used as the reference to normalize the transfection efficiency.

**Fig. S6.**
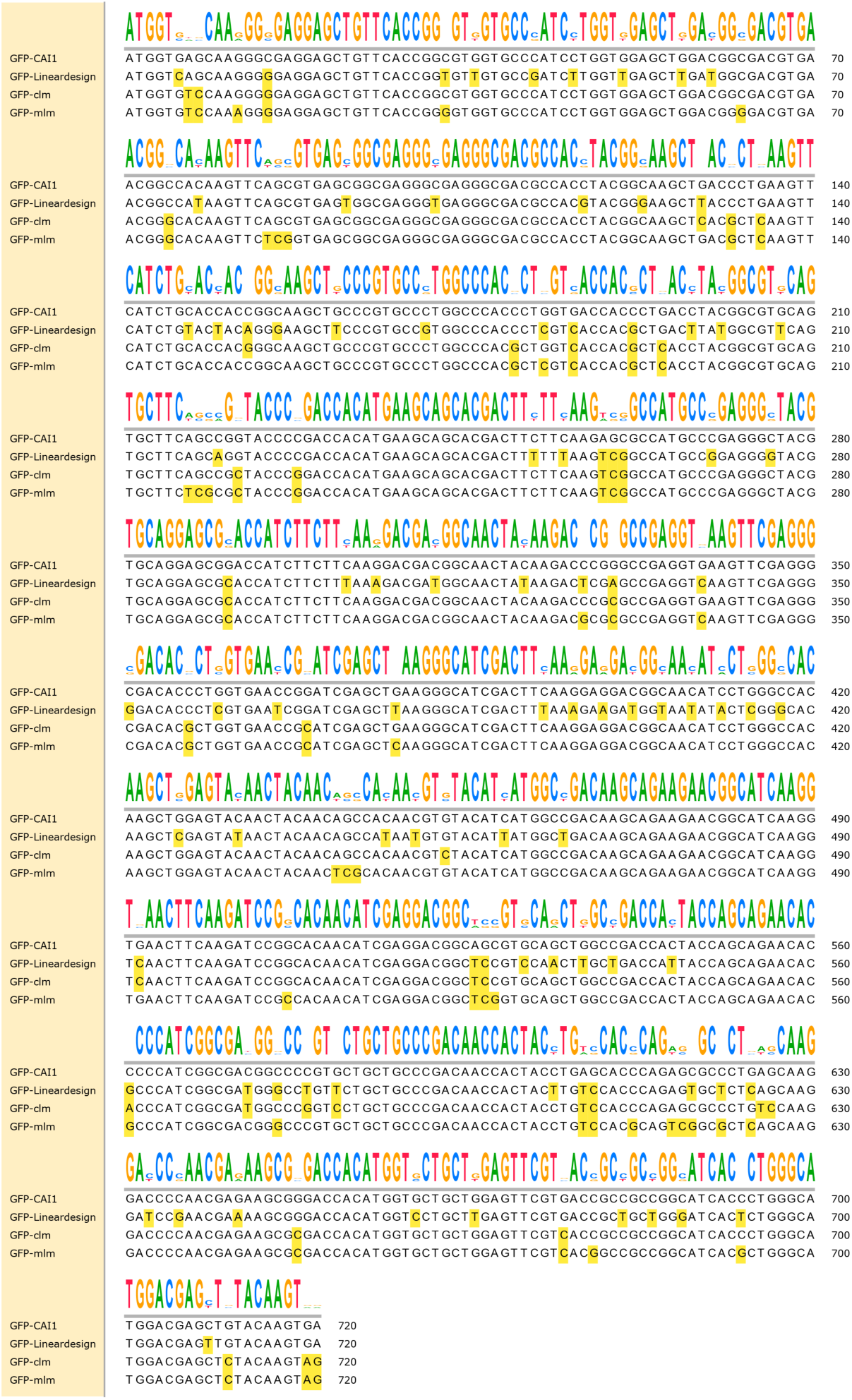
The multiple-sequence alignment of GFP candidates. The GFP candidates are obtained from CAI, Lineardesign, BiooBang (CLM) and BiooBang (MLM), respectively. The site variants are colored in yellow. Sequence logos derived from the multiple-sequence alignment.

**Fig. S7.**
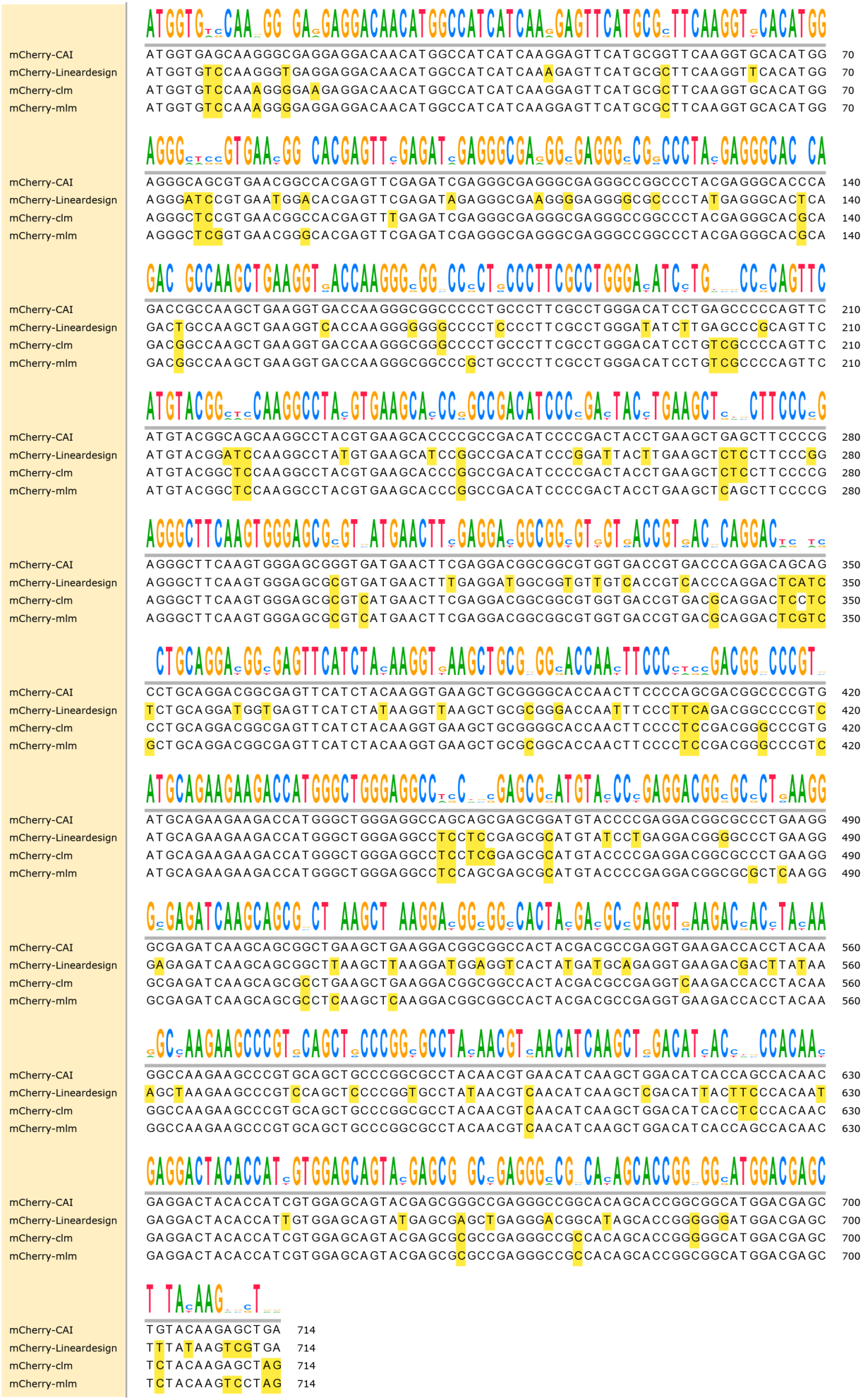
The multiple-sequence alignment of mCherry candidates. The mCherry candidates are obtained from CAI, Lineardesign, BiooBang (CLM) and BiooBang (MLM), respectively. The site variants are colored in yellow. Sequence logos derived from the multiple-sequence alignment.

**Table S1.** Detailed language model comparison on the protein-related tasks.

|  | BiooBang | Ankh | Ankh-<br>base | ProtT5-xl-U50 | ESM-1b | ESM2-33 | ESM2-36 |
| --- | --- | --- | --- | --- | --- | --- | --- |
| GB1P(regression) | **0.91** | 0.84 | <u>0.85</u> | 0.78 | 0.81 | 0.82 | 0.81 |
| FluP(regression) | **0.66** | <u>0.62</u> | 0.61 | 0.58 | 0.5 | 0.48 | 0.48 |
| SolP(classification) | <u>0.751</u> | **0.764** | 0.742 | 0.744 | 0.673 | 0.75 | 0.749 |
| LocP(classification) | <u>0.827</u> | **0.832** | 0.814 | 0.832 | 0.8 | 0.818 | 0.824 |
| FoldP(classification) | **0.691** | <u>0.611</u> | 0.588 | 0.576 | 0.576 | 0.563 | 0.605 |
| SSP(classification) | 0.760 | **0.785** | 0.769 | 0.768 | 0.744 | 0.774 | <u>0.776</u> |

**Table S2.** Detailed language model comparison on the mRNA-related tasks.

|  | BiooBang | CodonBert | CaLM |
| --- | --- | --- | --- |
| VDP(regression) | 0.8266 | 0.7046 | <u>0.8071</u> |
| RFPE(regression) | 0.8382 | <u>0.8314</u> | 0.7427 |
| FEP(regression) | 0.7746 | 0.7692 | <u>0.7696</u> |
| EEP(classification) | 0.545 | 0.468 | <u>0.525</u> |

**Table S3.** Details of the 5-fold cross-validation to predict LnTE.

|  |  | fold1 | fold2 | fold3 | fold4 | fold5 | std | mean |
| --- | --- | --- | --- | --- | --- | --- | --- | --- |
| BiooBang-CDS | Muscle | 0.5769/<br>0.8936 | 0.6835/<br>0.8235 | 0.5287/<br>0.9293 | 0.3411/<br>0.9554 | 0.5650/<br>0.7915 | 0.1248/<br>0.0695 | 0.5390/<br>0.8787 |
|  | HEK | 0.5398/<br>0.6763 | 0.5508/<br>0.6753 | 0.4875/<br>0.7183 | 0.5599/<br>0.6787 | 0.5851/<br>0.6094 | 0.0360/<br>0.0392 | 0.5446/<br>0.6716 |
|  | PC3 | 0.6228/<br>0.5825 | 0.6224/<br>0.5610 | 0.6037/<br>0.5898 | 0.6119/<br>0.5791 | 0.6004/<br>0.5791 | 0.0103/<br>0.0106 | 0.6122/<br>0.5783 |
| BiooBang-UTR | Muscle | 0.5618/<br>0.8709 | 0.4825/<br>0.8066 | 0.4992/<br>0.7525 | 0.4850/<br>0.7535 | 0.4363/<br>1.033 | 0.0452/<br>0.1166 | 0.4930/<br>0.8433 |
|  | HEK | 0.4223/<br>0.772 | 0.4174/<br>0.7642 | 0.4235/<br>0.7573 | 0.4015/<br>0.7563 | 0.4350/<br>0.7622 | 0.0122/<br>0.0063 | 0.4199/<br>0.7624 |
|  | PC3 | 0.4787/<br>0.6504 | 0.4732/<br>0.6616 | 0.4848/<br>0.6719 | 0.5050/<br>0.6729 | 0.4228/<br>0.6919 | 0.0305/<br>0.0154 | 0.4729/<br>0.6697 |
| UTR-LM | Muscle | 0.3812/<br>1.1374 | 0.3920/<br>1.0982 | 0.3282/<br>1.1214 | 0.4215/<br>0.9525 | 0.4962/<br>1.0286 | 0.0617/<br>0.0766 | 0.4038/<br>1.0676 |
|  | HEK | 0.3983/<br>0.7417 | 0.3530/<br>0.7716 | 0.4004/<br>0.7520 | 0.3990/<br>0.7201 | 0.3944/<br>0.7204 | 0.0203/<br>0.0219 | 0.3890/<br>0.7411 |
|  | PC3 | 0.4241/<br>0.6536 | 0.4245/<br>0.6498 | 0.4449/<br>0.6574 | 0.4482/<br>0.6601 | 0.4209/<br>0.6800 | 0.0130/<br>0.0118 | 0.4325/<br>0.6602 |

**Table S4.** Details in predicting MRL on the independent test set.

|  | UTR-Human | UTR-Random |
| --- | --- | --- |
| BiooBang-UTR | 0.856/0.111 | 0.934/0.113 |
| UTR-LM | 0.801/0.517 | 0.870/0.548 |

**Table S5.** Detail of the ablation study on two fine-tuning strategies with frozen or full parameter.

|  | Frozen params | Full params |
| --- | --- | --- |
| BiooBang-UTR | 0.6619 | 0.9652 |
| UTR-LM | 0.6 | 0.962 |

